# CMV can spread through plant to plant contact: implications for experimental practices

**DOI:** 10.1101/2025.07.30.667662

**Authors:** Lola Chateau, Marion Szadkowski, Jérémy Théodore, Loup Rimbaud

## Abstract

Cucumber mosaic virus (CMV) is a major plant pathogen with a worldwide distribution and the widest host range among all known plant viruses. It affects numerous crop species and can cause symptoms that significantly reduce yield. CMV is primarily transmitted by aphids and more sporadically through seeds. It is frequently studied in laboratory settings with the aim of developing effective control strategies. In many experiments, infected plants are placed in direct contact with healthy ones assuming that CMV cannot be transmitted in this way. However, this has not been formally demonstrated. Therefore, this study aimed to assess whether CMV can be transmitted through plant-to-plant contact. Infected plants were first rubbed against healthy ones and then left in contact for 28 days. Target plants were subsequently tested using DAS-ELISA to detect potential transmission. We applied this protocol in two separate experiments totalizing 15 combinations of plant species including pepper (*Capsicum annuum*) and five weed species commonly found in Espelette pepper fields (*Capsella bursa*-*pastoris, Cerastium glomeratum, Stellaria media, Stachys arvensis* and *Trifolium repens*). We found that CMV could be transmitted through contact between pepper and all tested weed species except *T. repens*. These findings highlight the importance of verifying whether a virus is capable of contact transmission before carrying out experiments in conditions that could lead to such contacts. In case of transmission, appropriate precautions will be crucial to avoid unintended transmissions.

## Introduction

Plant-to-plant transmission of several plant viruses, such as potato virus Y or zucchini yellow mosaic virus, is known to be possible by simple contacts between plants, although their primary mode of transmission is aphid vection (Coutts, Kehoe, and Jones 2013; Coutts and Jones 2015). Strict measures must be implemented during experiments involving those viruses to prevent accidental contamination. Indeed, experiments in plant virology often involve a large number of plants. Whether conducted in fields, greenhouses, or trays, these plants may come into physical contact due to their close proximity in experimental facilities. Environmental perturbations such as wind or ventilation can further promote friction between neighbouring plants. Handling plants, such as touching infected plants before healthy ones, or using watering hoses that come into contact with both infected and healthy plants, can also represent a potential source of contact transmission. Experimental protocols involving these viruses are designed to avoid plant-to-plant contact. However, such precautions may not be taken with viruses for which no contact transmission has been reported. Since data on the topic are scarce, it is essential to test whether a virus can be transmitted through plant-to-plant contact before conducting experiments involving many plants in a confined space.

This is particularly important when assessing viral transmission by aphids, for which inadvertent contact transmission would compromise the results. There are two main approaches to measuring the aphid-mediated transmission of a virus. The first one is in completely controlled conditions, and consist in placing one or several aphids on an infected ‘source’ leaf and then transferring the aphids to the leaf of a healthy ‘target’ plant. After a certain period, target plants are diagnosed and the proportion of infected ones gives the efficiency of transmission (Doumayrou et al. 2013; Lehrer et al. 2007). The second approach comes closer to natural transmission and relies on arenas. In this setup, aphids are placed in a closed environment, containing both infected source plant and healthy targets. The proportion of infected target plants gives the propensity of transmission (Chatzivassiliou et al. 2016; Kanavaki et al. 2006). Aphids should be winged to better represent natural conditions of viral transmission, and effectively move between source and target plants within the arena experiments (Yuan 1996). However, managing winged aphids is challenging due to the difficulty of synchronizing generations in aphid breeding facilities, as well as the risk of escape beyond the experimental arenas. As a result, non-winged (apterous) aphids are often used instead (Katis et al. 2006; Claflin, Thaler, and Power 2015; Webb and Leng Kok-Yokomi 1993; Hobbs et al. 2000). To allow their movement between plants, the plants are positioned in direct contact with one another, which could inadvertently enable contact transmission of the virus, should this mechanism be possible. This would lead to inaccurately calculate transmission rates.

As mentioned before, for viruses that do not primarily rely on plant-to-plant contact for transmission, their ability to spread in this way is not always tested, and experimental setups are not necessarily designed with this eventuality in mind. This is the case of cucumber mosaic virus (CMV), a virus belonging to the genus *Cucumovirus* within the *Bromoviridae* family. CMV is present worldwide and has the largest host range among plant viruses (Hirsch and Moury 2021). It is primarily transmitted by aphids, with more than 80 species known to act as vectors. It is also transmitted by the seed of some of its host plants. It induces various symptoms in plants, including mosaic patterns, yellowing, and leaf and fruit distortions, among others (Hirsch and Moury 2021). It is the most economically important virus for six different crop varieties and the first virus affecting annual crops in 14 countries (Gallitelli 2000). For these reasons, CMV is well studied in laboratory, and even serves as a model for studying plant viruses (Scholthof et al. 2011).

In Espelette pepper, the virus induces crop losses not only in quantity (yield) but also in quality (Lepage et al. 2025). Implementing efficient management strategies requires a fine understanding of the epidemic dynamics in the local context. This supposes for instance measuring aphid-mediated transmission from and to plants commonly found in the Espelette landscape. However, to the best of our knowledge, the possibility of plant-to-plant contact transmission of CMV is not documented, it is thus crucial to test it. This study evaluates whether precautions should be taken in experiments to avoid inadvertent transmissions.

## Material and methods

The test has been carried out in two independent experiments.

### Virus material, plant material, and greenhouse conditions

#### Virus isolate

The CMV isolate used in this study was collected on an infected pepper (*Capsicum annuum*, cultivar ‘Gorria’) in the area of Espelette in 2022 (Gaudin et al. 2025). It belongs to the taxonomic subgroup IB of CMV and was stored in dried leaves (Bos 1983) from tobacco plants (*Nicotiana tabacum*, cultivar ‘xanthi’) and conserved at 4°C.

#### Plant material

The following plant species, commonly found in the Espelette landscape, were included in the contact transmission experiments: Espelette pepper (*Capsicum annuum* cultivar ‘Gorria’, seeds collected from farmers, who practice mass selection), clammy chickweed (*Cerastium glomeratum*, seeds collected in Espelette pepper fields), staggerweed (*Stachys arvensis*, seeds collected in Espelette pepper fields), shepherd’s purse (*Capsella bursa*-*pastoris*, seeds produced in our experimental facilities), chickweed (*Stellaria media*, commercial seeds from “Arbiotech”), and white clover (*Trifolium repens*, commercial seeds from “Semences du Puy”).

#### Greenhouse conditions

All plants were grown in an insect-proof greenhouse from October to December (first experiment) and from April to June (second experiment) in Avignon, France. Greenhouse temperatures ranged from 11.9°C to 25.1°C during the first experiment and 11.4°C to 37.9°C during the second experiment.

### Mechanical inoculation and virus detection

Mechanical inoculations and viral detection both require the preparation of plant homogenates. For this, one gram of plant material (dried or fresh leaves) was ground in 4 mL of extraction buffer containing 0.03M sodium phosphate (Na_2_HPO_4_) and 0.2% diethyldithiocarbamate (DIECA).

For mechanical inoculations, every homogenate was mixed with 90 mg of carborundum and 90 mg of activated charcoal before being applied to 2 leaves per plant using two strokes. After five minutes, the leaves were rinsed with tap water.

For virus detection, a DAS-ELISA was performed on the homogenates following the method described by (Hirsch et al. 2023). Plants were considered positive when their optical density (OD) at 405 nm was higher than three times the OD of a healthy control, and negative otherwise. Symptom assessment was also conducted, focusing on the presence of mosaic, deformation and necrosis on leaves.

### Mechanical inoculations to prepare source plants

To obtain CMV infected plants serving as sources for contact transmission experiments, tobacco plants (*N. tabacum* and *N. benthamiana*) were first mechanically inoculated with CMV from dried material, and maintained in a greenhouse for 14 days post-inoculation (dpi). Then, a homogenate of these infected tobacco plants was used to mechanically inoculate 5-week-old plants of the 6 tested species: *C. annuum, C. bursa-pastoris, C. glomeratum, S. media, S. arvensis* and *T. repens*. Ten (first experiment) or twelve (second experiment) plants per species were inoculated, except for *C. annuum* in the second experiment (17 plants), and *T. repens* for which 100 (first experiment) or 105 plants (second experiment) were inoculated as infection was difficult to achieve in preliminary experiments (data not shown). All these plants were diagnosed by DAS-ELISA at 28 dpi. Source plants in the contact transmission experiments were randomly chosen among the infected plants.

### Contact transmission experiments

Contact transmission was tested in two independent experiments. Each experiment included several transmission combinations: from *C. annuum* to *C. annuum*, from *C. annuum* to the five weed species and from each weed species to *C. annuum*. In the second experiment, additional combinations were tested: transmission from each weed species to itself, except for *T. repens* for which not enough infected plants had been obtained. This resulted in a total of 15 different combinations (listed in Table 1). For each combination, one infected source plant was put into contact with eight 5-week-old healthy target plants. Each combination was tested once per experiment, except for the *C. annuum* to *C. annuum* combination in the second experiment, which was replicated four times. In every experiment, 12 healthy plants served as negative controls (two per species).

**Table 1.** Number of infected and symptomatic plants in the contact transmission experiments. Symptomatic plants were identified based on the presence of typical CMV symptoms, including mosaic, deformation, and necrosis. Infected plants were diagnosed based on DAS-ELISA tests. Combinations resulting in contact transmission are highlighted in bold.

| Source plant | Target plants | Number of symptomatic plants |  | Number of infected plants |  | Transmission rate |
| --- | --- | --- | --- | --- | --- | --- |
|  |  | Exp. 1 | Exp. 2 | Exp. 1 | Exp. 2 |  |
| *Capsella bursa-pastoris* | *Capsicum annuum* | 4/8<sup>a</sup> | 3/8 | **5/8** | **7/8** | **75%** |
| *Cerastium glomeratum* | *Capsicum annuum* | 1/8 | 1/8 | **1/8** | **1/8** | **13%** |
| *Stellaria media* | *Capsicum annuum* | 1/8 | 2/8<sup>a</sup> | **1/8** | **1/8** | **13%** |
| *Trifolium repens* | *Capsicum annuum* | 0/8 | 0/8 | 0/8 | 0/8 | 0% |
| *Stachys arvensis* | *Capsicum annuum* | 5/8<sup>a</sup> | 1/8 | **6/8** | **1/8** | **44%** |
| *Capsicum annuum* | *Capsicum annuum* | 0/8 | 1/8 | 0/8 | **1/8** | **5%** |
|  |  |  | 1/8 |  | **1/8** |  |
|  |  |  | 0/8 |  | 0/8 |  |
|  |  |  | 0/8 |  | 0/8 |  |
| *Capsicum annuum* | *Capsella bursa-pastoris* | 0/8 | 0/8 | **5/8** | **4/8** | **56%** |
| *Capsicum annuum* | *Cerastium glomeratum* | 0/8 | 0/8 | 0/8 | 0/8 | 0% |
| *Capsicum annuum* | *Stellaria media* | 0/8 | 2/8<sup>b</sup> | 0/8 | 0/8 | 0% |
| *Capsicum annuum* | *Trifolium repens* | 0/8 | 0/8 | 0/8 | 0/8 | 0% |
| *Capsicum annuum* | *Stachys arvensis* | 0/8 | 0/8 | **1/8** | **1/8** | **13%** |
| *Capsella bursa-pastoris* | *Capsella bursa-pastoris* | NA | 0/8 | NA | **8/8** | **100%** |
| *Cerastium glomeratum* | *Cerastium glomeratum* | NA | 2/8 | NA | **2/8** | **25%** |
| *Stellaria media* | *Stellaria media* | NA | 0/8 | NA | **2/8** | **25%** |
| *Stachys arvensis* | *Stachys arvensis* | NA | 0/8 | NA | **7/8** | **88%** |

<sup>a</sup> 1 plant exhibited symptoms but was tested negative by DAS-ELISA.
<sup>b</sup> 2 plants exhibited symptoms but were tested negative by DAS-ELISA.
NA: Combinations not tested during the first experiment.

To perform the transmission test mimicking situations where plants might come into contact and potentially transmit viruses, each CMV-infected source plant was gently rubbed against the 8 target plants for 10 seconds, then placed at the centre of the group of target plants, allowing them grow while remaining in direct contact, a process known as “intertwining”. The experimental setup is illustrated in Figure 1. Plants were then kept in an aphid free greenhouse for 28 days. At the end of this period, each target plant was tested using DAS-ELISA to determine whether contact transmission had occurred.

**Figure 1.**
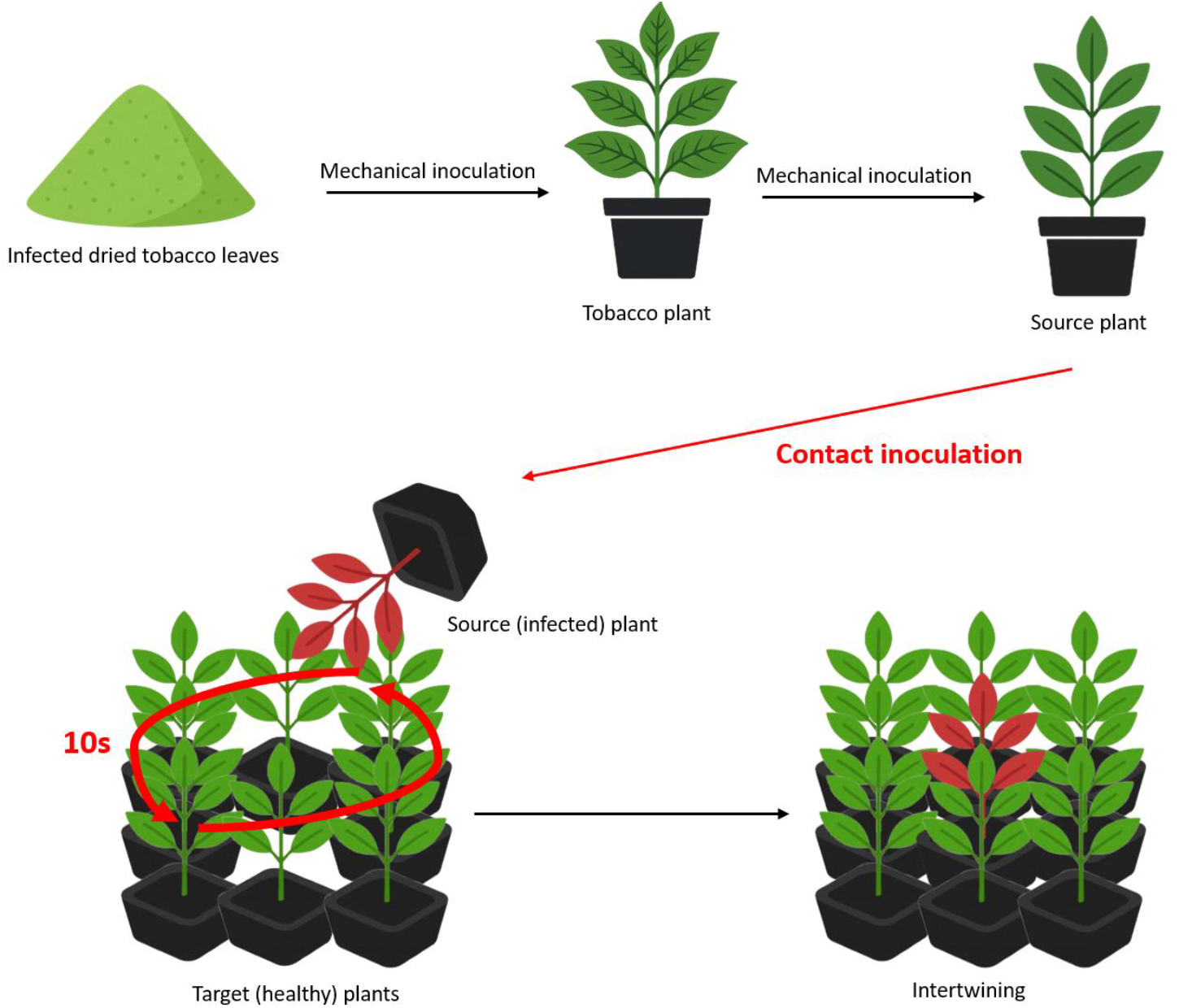
Scheme of the experimental setup to test for viral contact transmission. Infected source plants are obtained after successive mechanical inoculations. Then, a single source plant is rubbed against 8 healthy target plants during 10 seconds and next placed in the middle for 28 days of intertwining.

## Results

### Mechanical inoculations to prepare source plants

To obtain infected source plants, mechanical inoculations were performed using dried leaves from infected tobacco plants as inoculum. The success rate of mechanical inoculation (based on DAS-ELISA) varied by species (Figure 2 & Table S1), reaching 100% for *C. annuum* and *C. bursa-pastoris*, 90–100% for *S. arvensis*, 80–100% for *S. media*, 70–75% for *C. glomeratum*, and only 0.9–4% for *T. repens*. Symptom assessment showed consistent expression in *C. annuum* (90–94%), while results fluctuated between tests for the weeds: *C. bursa-pastoris* (0–50%), *C. glomeratum* (0–75%), *S. media* (70–92%), and *S. arvensis* (40– 100%). No symptom assessment was conducted for *T. repens* due to the difficulty of identifying CMV symptoms on this species.

**Figure 2.**
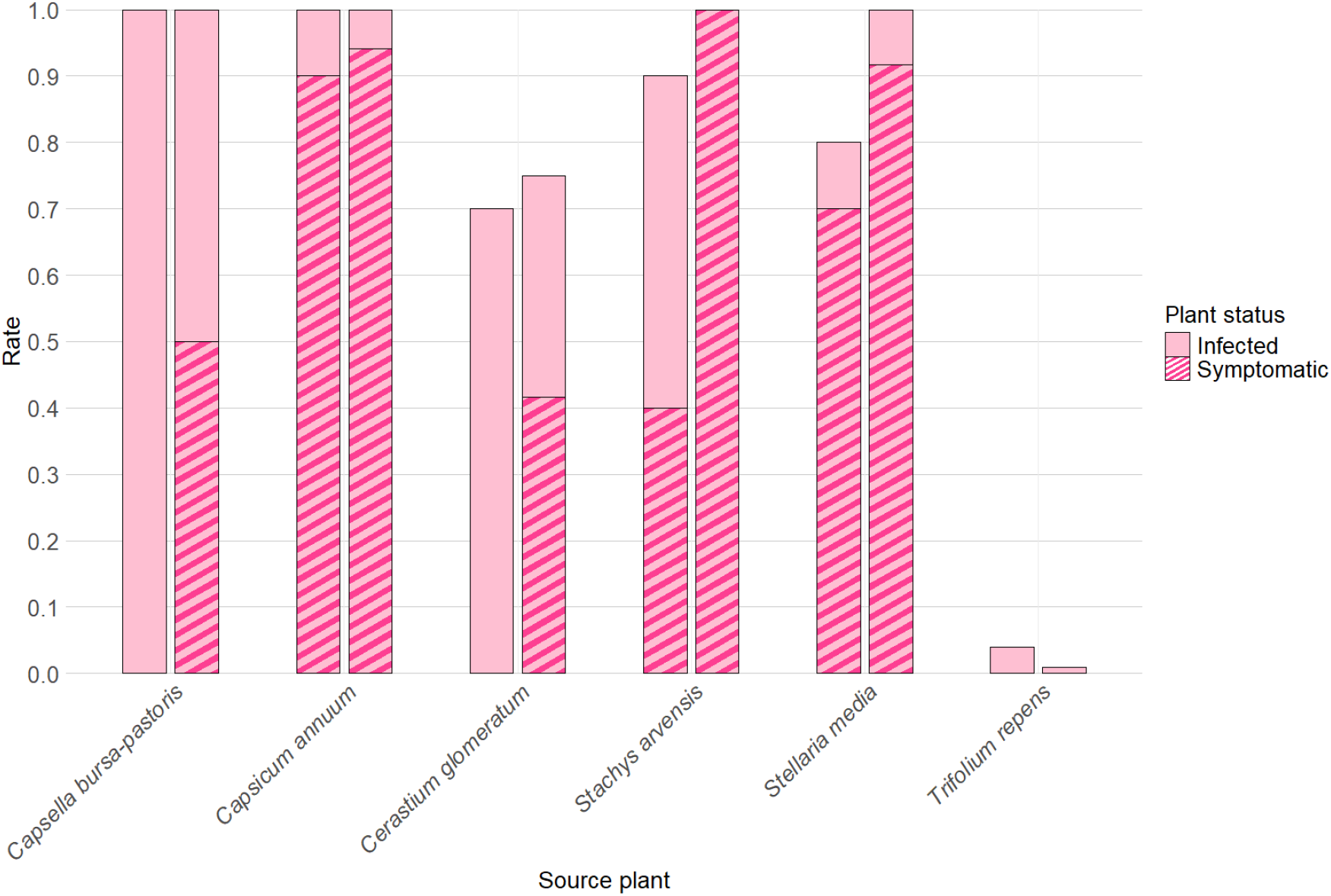
Rates of mechanical transmission. Bars represent infected plants (pink), including symptomatic ones (striped) for each plant species. The first bar corresponds to experiment 1 (n=10 plants except for *T. repens* (n=100 plants)), the second to experiment 2 (n=12 plants except for *C. annuum* (n=17) and *T. repens* (n=105 plants))

### Contact transmission experiments

Contact transmission was tested by rubbing infected and healthy plants and allowing them to grow in direct contact. Each target plant was then monitored for symptom expression and its sanitary status (healthy or infected) checked by DAS-ELISA (Table 1).

Transmission through contact was detected in 11 out of the 15 plant combinations tested. Transmission to *C. annuum* was observed from *C. bursa-pastoris* (75%), *C. glomeratum* (13%), *S. media* (13%), and *S. arvensis* (44%). In the opposite direction, *C. annuum* transmitted CMV to *C. bursa-pastoris* (56%) and to *S. arvensis* (13%). Transmission also occurred between *C. annuum* plants, although at a very low rate (5%). Transmission between weeds of the same species was also observed: 100% for *C. bursa-pastoris*, 25% for *C. glomeratum*, 25% for *S. media*, and 88% for *S. arvensis*. No transmission was detected from or to *T. repens*. Likewise, no infection was recorded from *C. annuum* to *C. glomeratum* or to *S. media*. Self-transmission in *T. repens* could not be tested due to the limited number of infected source plants.

Among the 20 peppers displaying symptoms, only 17 tested positive by DAS-ELISA. This represents a specificity of symptom monitoring of 85% (number symptomatic plants being infected/number of symptomatic plants). In contrast, 8 peppers tested positive for CMV without exhibiting any symptom. Symptoms on weed plants were detected only on *S. media* and *C. glomeratum* in the second experiment with a frequency of 25%.

None of the negative control plants presented symptoms nor was tested positive in DAS-ELISA (data not shown)

## Discussion

This study emphasizes the importance of verifying whether a virus can be transmitted through contact before conducting experiments with plants in close proximity. Indeed, the absence of prior reports does not necessarily mean that such transmission is impossible. For aphid arena experiments, such verification can easily be achieved by including a control combination without aphids to confirm that contact transmission does not occur between the tested plants. With respect to CMV, the results demonstrate that contact transmission can occur between certain plant combinations, with all tested species being able to be source or target, except *T. repens*. This conclusion, without being generalizable to other species, demonstrates the need for caution and the need to minimize plant contacts during experiments to avoid biases in the results. Indeed, slight friction between plants can occur spontaneously, even without direct intervention, due to wind or air circulation. Should micro-wounds be caused to the leaves, for example due to airborne sand particles, the virus could be transmitted. One potential solution is to increase the distance between plants in greenhouses whenever possible. To avoid plant contact in aphid arena experiments with apterous aphids, artificial bridges could allow aphid movement between plants.

Mechanical inoculation does not exhibit equal efficiency across all plant species. Transmission rates ranged from 100% in *C. annuum* and *C. bursa-pastoris* to only 4% maximum, in *T. repens*. Although mechanical inoculation of CMV is generally very effective (Jacquemond 2012), the low success rate in *T. repens* required inoculating up to 105 plants, compared to just a dozen for the other species. This may explain the absence of contact transmission involving *T. repens* in our experiments. Another factor that could explain the varying success of both mechanical and contact transmission of CMV between different species is the origin of the isolate used. Only one CMV isolate, originating from an infected *C. annuum* plant, was used in the experiments. Although CMV has a broad host range, this particular isolate may be better adapted to pepper than to some of the weed species.

During symptom assessment, some asymptomatic plants tested positive by DAS-ELISA, suggesting two possible explanations. First, these plants could be still in the viral incubation period. Although the plants had been in contact for a month at the time of symptom evaluation, the exact moment of transmission remains unknown. It could have occurred early on during the leaf rubbing process or later during the course of the experiment through plant intertwining. If the transmission happened only a few days before the experiment ended, the virus may have started replicating but not yet progressed enough to cause visible symptoms. The second possibility is that some plants will never show any symptoms, as suggested by the outcome of the mechanical inoculations (Figure 1). Such variability in symptom expression within species can be a consequence of the genetic variability of the seeds used, in particular those coming from the field (*C. glomeratum* and *S. arvensis*). Additionally, the two experiments were conducted at different times of the year, leading to variations in temperature and light, which could affect symptom expression in spite of our efforts to standardize the protocol.

At the opposite, some plants recorded as symptomatic were actually negative in DAS-ELISA. This could be due to the lack of specificity in our visual assessment, meaning that some leaves had been incorrectly interpreted as CMV symptoms. Another possibility is that the DAS-ELISA test lacked sufficient sensitivity to detect very low viral loads in the plant or that the virus had a heterogeneous accumulation within the plant.

The transmission protocol applied in this study combined two techniques: leaf rubbing, where plants were physically rubbed against each other, and intertwining, where plants were left to grow in contact. This protocol was destined to maximize the possible contacts between plants. In case of absence of contact transmission, experiments with plants in close proximity (such as in arena tests) can then be performed without any risk. Otherwise, plants should be spaced to avoid inadvertent transmission. The exact mechanism underlying contact transmission of CMV remains unclear. Additional experiments could provide more detailed insights into these mechanisms, such as testing transmission with only one of the two techniques done here. Another approach could involve applying drops of viral homogenate directly onto leaves to assess whether this is sufficient to cause infection. Nevertheless, these investigations lie beyond the scope of the present study.

Our experimental setup does not necessarily reflect the real contact between plants in the field, leaving the impact of contact transmission unclear with respect to CMV epidemiology. Nevertheless, transmission between pepper plants appears rare: no transmission was observed in the first experiment, and only one plant in two of the four replicates became infected during the second. This suggests that contact transmission in peppers is infrequent, which is reassuring for both past and future experiments involving peppers, as well as for farmers growing peppers in close proximity in the field.

## Authors contributions

LC and LR conceived and designed the experiments, and analyzed the data. LR, LC, MS, and JT carried out the experiments. LC, MS, and LR wrote the manuscript. LR supervised the project and secured funding.

## Acknowledgements

We thank the staff of the Experimental facilities of the PROPHYLE platform from the Plant Pathology research unit (https://doi.org/10.15454/8DGF-QF70) who ensured the production and maintenance of the plants and plant-growth facilities that allowed us to carry out this work. We also thank Alice Conilh and Lucas Gonzalez for their experiments that inspired the present work, as well as Benoît Moury, Cécile Desbiez and Pauline Ezanno for stimulating discussions. We thank Lia Lamacque and Cécile Desbiez for constructive comments on earlier version of this manuscrit. Finally, we thank Damien Richard for his interest and recommendation of this article in PCI Infections (https://doi.org/10.24072/pci.infections.100255; Richard, 2025), based on the insightful peer-reviews by Cica Urbino, Véronique Brault and one anonymous reviewer.

## Data Availability

All data relevant to this work are available within the article and in Tables S1 and S2.

## Supporting information

**Table S1**. Number of symptomatic (based on visual monitoring) and infected (based on DAS-ELISA diagnostic) plants after mechanical inoculation of CMV to 6 plant species. No symptom assessment was conducted for T. repens due to the difficulty of identifying CMV symptoms on this species.

**Table S2**. Optical density (OD) values from DAS-ELISA. The ‘Experiment’ column indicates whether the data come from the first or second experiment. The ‘Plant’ column specifies whether the plant was a source plant, assessed after mechanical transmission or a target plant, assessed after contact transmission. The ‘Combination’ column indicates the plant species; for target plants the leftpart of the arrow indicates the source of the transmission. The ‘Number’ column indicates the index of the plant. OD was measured between 3 h and 4 h of incubation of the DAS-ELISA substrate.

