## Supplementary material for "CMV can spread through plant to plant contact: implications for experimental practices": Table S1

**Table S1**. Number of symptomatic (based on visual monitoring) and infected (based on DAS-ELISA diagnostic) plants after mechanical inoculation of CMV to 6 plant species. No symptom assessment was conducted for *T. repens* due to the difficulty of identifying CMV symptoms on this species.

| ***Species*** | **Experiment 1** | | **Experiment 2** | |
|---|---|---|---|---|
| | **Symptomatic / inoculated** | **Infected / inoculated** | **Symptomatic / inoculated** | **Infected / inoculated** |
| *Capsicum annuum* | 9/10 | 10/10 | 16/17 | 17/17 |
| *Capsella bursa-pastoris* | 0/10 | 10/10 | 6/12 | 12/12 |
| *Stachys arvensis* | 4/10 | 9/10 | 12/12 | 12/12 |
| *Stellaria media* | 7/10 | 8/10 | 11/12 | 12/12 |
| *Cerastium glomeratum* | 0/12 | 7/10 | 5/12 | 9/12 |
| *Trifolium repens* | - | 4/100 | - | 1/105 |
